# SUSTAINABLE MICROBIAL BIOTRANSFORMATION OF CR(VI) TO CR(III) IN TANNERY EFFLUENT AND ITS VALORIZATION INTO CR(III) NANOPARTICLES VIA *TRIDAX PROCUMBENS*-MEDIATED GREEN SYNTHESIS

**DOI:** 10.64898/2026.04.23.720289

**Authors:** Neethu Asokan

## Abstract

Environmental pollution from leather industries have become a menace. The microbial remediation of industrial waste and it’s reuse for agriculture could be a beneficial outcome. In present study, the bioremediated Cr III in the effluents are further converted to value product – Chromium oxide NP. This ensures double edged benefit as effluent is bioremediated and Chromium oxide NP with several applications is derived. A noteworthy advancement of the research involved the green synthesis of chromium oxide nanoparticles using Tridax procumbens. The effluent bioremediated can be used for agricultural purposes. By effectively characterizing tannery effluent and isolating chromium-tolerant bacteria, the study not only demonstrate a practical bioremediation solution but also showcase the potential of green synthesis in producing chromium oxide nanoparticles. In conclusion, this research marks a significant advancement in environmental science, leveraging both biological and nanotechnological innovations to address pressing challenges in pollution control. The present study focuses on a novel process of obtaining chromium oxide nanoparticle from tannery effluent with several applications derived from bioremediated tannery effluent using a cost-effective and eco-friendly process. The nanoparticle has a stable particle size and exhibit antioxidant, anti-diabetic properties. This product offers a breakthrough solution for the leather industry and healthcare sector.

**GRAPHICAL ABSTRACT:** 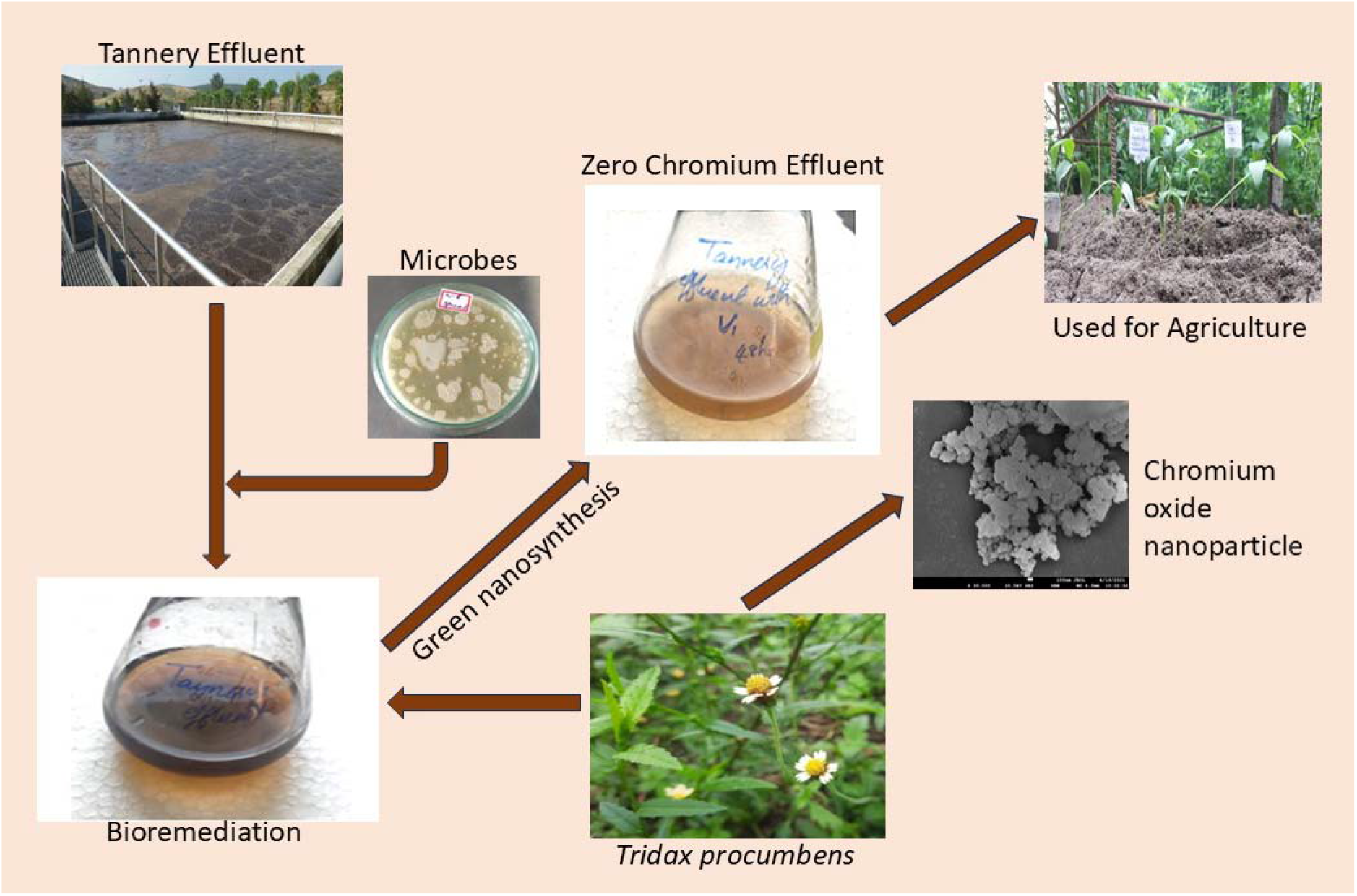

## 1. INTRODUCTION

Bioremediation is the biological degradation of organic waste under precise conditions. Bioremediation can also be defined as the removal of toxic pollutant from environment or atmosphere through biological system without the production of any secondary pollutant. It has been of advance research for its cost effectiveness and efficiency in metal treatment (Patil et al., 2024). Bioremediation technology using microorganisms was invented by George M. Robinson. It is the use of living organisms, primarily microorganisms, bacteria and fungi, to degrade the environmental contaminants into less toxic forms (Wang et al., 2000; Xia et al., 2019). Microorganisms can be isolated from any environmental conditions (Campo et al., 2019; Cervantes et al., 2007). As a result of the adaptability of microbes and additional biological systems, these can be used to degrade or remediate environmental menaces. Bioremediation mechanisms may include bioaugmentation, biostimulation, biotransformation, bioprecipitation, biocrystallization and bioleaching (Mahamuni-Badiger et al., 2023). There are reports stating that bacteria utilize mechanisms like biosorption and bioaccumulation to remediate the contaminants through redox reactions and enzyme transformation methods (Patil et al., 2024). Environmental heavy metal pollution by anthropogenic and industrial activities has triggered substantial damage to the ecosystems (Ali et al., 2023; Chai et al., 2018). The heavy metal sources include the mining and smelting of ores, leather industry effluent, effluent from storage batteries, or from the manufacturing and inadequate use of fertilizers, pesticides. There are several previous reports of bioremediation of Chromium contaminated source with the application of microbes (Prusty et al., 2019; Gadd, 2004; Lloyd, 2002). The demand for leather has been accelerating past some years and the increasing global population the leather industries flourished. Tanning is a vital process for converting raw hides and skins into finished leather. Chrome tanning is the most used process and this contributes to severe heavy metal contamination of environment (Zuriaga-Agustí et al., 2015). Indian leather industry is of pre-eminence in the global economy. India houses for about 2000 tanneries. The maximum collection of tanneries in India is on the banks of the Ganga River in North India and the Palar River in Tamil Nadu. India is the second largest leather exporter in the world. Tamil Nadu alone has about 750 operational units, 650 leather garment units and 497 leather shoe units (**Central Leather Research Institute, Chennai, 1997**). According to Central Leather Research Institute and the State Pollution Control Board, Chennai, the figure of tanneries situated in Vellore District has more tanneries in Ranipet (41.61 per cent); followed by Vaniyambadi (25.18 6 per cent). The tanning activity demands proper supply of water. The majority of the tanneries are hence situated around the bank of Palar River near Tamil Nadu. The tanneries have thus become a pollution concern when the tanneries were shifted from Vegetable Tanning to Chrome Tanning in the early seventies since that is a time saving activity. Leather Industries releases about 3,000–3,200 liters of water per 100 kg of skins and hides treated. The effluent is released as waste water, remaining confluent in the water body. The conventional nature of tanning activity leads to the discharge of malodorous bearing effluent, and toxic compounds. It comprises organic constituents in the form of fatty acid, protein, and inorganic components namely chlorides, trivalent chromium, nitrates, phosphorus, sulfides, and sulfates. The disposed effluent in the waters undesirable affects the aquatic life and is dangerous for the aquatic ecosystem. It also affects the physical, chemical, and biological character of waters and steals dissolved oxygen (DO) from waters. These wastes also cause damage to the soil quality, crop pattern, vegetation, and yield and affect the plants too (Varadaraj et al., 1997). Even though some large scale industries have treatment plan, most of the operational industries are small scale which do not have enough finance to set up a treatment plan.

Wastes are found to originate from all steps of leather making, such as fine leather particles, residues from various chemical discharges and reagents from different waste liquors including of large pieces of leather cuttings, trimmings and gross shavings, fleshing residues, solid hair debris and remnants of paper bags (Bhaskaran, 1977). From 1000 kg crude hide that goes into use, an average of 850 kg is generated as hard waste while tanning leathers. The industries generate ~ 30–35 L waste per kg skin/hides treated. And more than 80 per cent organic pollution loading in BOD are being generated from the beam house (pre-tanning); high from rotten hide and hair material. During tanning 300 kg chemical (lime, salt) are added per ton crude hides. Extra amount of non-used salts are seen in the waste waters along with chromates. Tannery wastes and wastes in solids enter surface waters, where the poisons are washed down stream and end up in waters that are for bathing, cooking, swimming, and irrigation. High levels of Cr, Ni, Zn, Pb are present in soil due to the chemicals use during tanning and settling of disposed wastes into ground and in soil (Manjunatha et al., 2011). Chromium wastes also enter into ground soils and end up in ground waters systems. The ability of Chromium to cause toxicity is widely recognized. The long term exposure to chromium can cause oxidative stress to the body and affects all major organs. Severe health effects including oxidative stress, DNA damage, respiratory disorers, kidney and liver dysfunction and lung cancers are some hazards due to chromium toxicity. Not all chromium is hazardous; however the chromium level in tannery effluents and the state of chromium is in fact fatal to plants, humans and animals (Brandon et al., 2024). One of the measures concentrated in previous study is the bioremediation using microbes and is believed to have boundless prospective for future development due to its environmental compatibility and cost-effectiveness (Mahdi et al., 2019; Campo and Salamarca, 2019). A wide range of microorganisms, including bacteria, fungi, yeasts, and algae, can be used biologically as active methylators, which are able to at least transform toxic species (Benazir et al., 2010; Focardi et al., 2013). *Bacillus sp* has been reported as major organism capable of bioremediation of hydrocarbons and heavy metals. They are known to have enzymes responsible for bioremediation of environment pollutants (Das et al., 2024). *Pseudomonas sp* and *Arthrobacter* is known to have chromate bioremediation properties; similarly with few fungal species like *Aspergillus niger* and *Penicillium chrysogenum* (Das et al., 2024). Many microbial detoxification procedures include the efflux or exclusion of metal ions from the cell, which may sometimes result in high local concentrations of metals at the cell surface, where they can react with biogenic ligands and precipitate (Kim, 1985). Even though microbes cannot destroy metals, they however can amend their properties through array of mechanisms. Hence the present project focuses on bio remediating Chromium in tannery effluent and its application in agriculture.

## 2. METHODOLOGY

### 2.1. Collection and characterisation of tannery effluent

The tannery effluent was collected in sterile bottles from local industries around Tirupattur District, Tamil Nadu, India and physio-chemical characteristics were analyzed.

### 2.2. Isolation of Cr (VI) reducing bacteria and effect of growth

The collected effluent was aseptically spread plated on to Nutrient agar plates with Cr (VI) at a concentration of 100µg/ml and incubated at 37 °C for 24–48 h. The effect of bacterial growth was determined in Minimal medium supplemented with Cr (VI) at different concentration (1-100 µg/ml) followed by incubation at 35 °C under shaking condition at 120 rpm. The tolerant isolate was selected for further study.

### 2.3. Bioremediation of Chromium using Chromium tolerant Microorganism

Tannery effluent was filtered using normal mesh filter and transferred to 250 ml flask to begin with a volume of 100 ml. The effluent was sterilized to remove other microorganisms. After sterilization potent organism isolated was inoculated into the flask and incubated at 37-degree Celsius for 24-48 hours and observed for remediation. The sample at interval of 24 hours was collected and estimated for chromium concentration and visible color changes due to bioremediation (Maurya et al., 2025).

### 2.4. Reduction of Cr (VI) in tannery effluent and analysis using diphenyl carbazide method

The Cr(VI) reduction potential of bacterial isolate was evaluated by growing the bacterium in Erlenmeyer flasks containing Cr(VI) effluent at different concentration and incubating at 35 °C under shaking condition at 120 rpm. The broth containing Cr(VI), but without bacterial culture was taken as control (Thacker et al., 2007). The bacterial growth was then monitored spectrophotometrically at 600nm. The Cr(VI) reduction potential was evaluated by a colorimetric method in terms of decrease in Cr(VI) concentration by using a Cr(VI) specific colorimetric S-diphenylcarbazide (DPC) method at 540 nm (APHA, 2012) (Thacker et al., 2007). The Cr(VI) reduction was calculated by using the following formula:

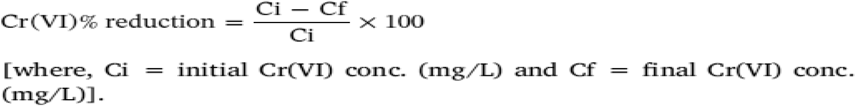

The results were analyzed statistically using appropriate statistical tools.

### 2.5. Molecular characterization of Cr (VI) reducing bacteria

The genomic DNA was prepared from overnight grown bacterial culture using the alkaline lysis method described by **Kapley et al. (2001)**. Approximately 5 μL DNA was used to amplify 16S rDNA gene using universal primers in a PCR thermocycler. The PCR product was analyzed on 1% agarose gel. The product was then sent for 16SrRNA sequencing to obtain the partial sequences (**Bhargava et al., 2018**). The partial sequences were subjected to BLAST analysis using the online source www.ncbi.nlm.nih.gov/BLAST that suggests the identity of isolated bacterium. The phylogenetic tree was constructed by neighbor-joining method using NCBI database online phylogenetic tree builder (http://www.ncbi.nlm.nih.gov). Further, the sequences were also deposited to Gene-Bank to obtain accession number.

### 2.6. Production of Chromium nanoparticles using *Tridax procumbens* (Green synthesis)

*Tridax procumbens* was collected from the college premises and washed thoroughly to remove sand and dirt. These were cut into small pieces and boiled in 100mL distilled water for 20 minutes. The filtrate was filtered using Whatmann paper and cooled. About 10 ml of the bioremediated tannery effluent was mixed with 10ml of *Tridax* extract. The solution was observed for a color change from orange to green indicating the formation of chromium oxide nanoparticles. The solid product was filtered and washed with ethanol and then dried at room temperature (Ramesh et al., 2012).

### 2.7. Characterisation of Chromium nanoparticles

The produced nanoparticles were characterized. X-ray powder diffraction was carried out for measuring the structure of nanoparticles. Transmission electron microscopy (TEM) was used to determine the size of chromium nanoparticles. Field emission Scanning electron microscopy with EDAX analyzer (Model no: JSM-7610F/X-MaxN) was used in this study for the elemental analysis of chemical characterization of samples. Zeta potential of chromium nanoparticle along with particle size was measured with particle analyzer.

### 2.8. Application of bio-remediated effluent for seed germination (Karunyal et al., 1994)

The treated and untreated waste/effluent was diluted and a control without effluent was maintained. The seeds were soaked in both treated and untreated effluent overnight and the germination of plants was studied up to 120 days. The Impact of these treatments on plant growth was measured. The size comparisons for internodes, shoot length, root length, total chlorophyll, and yield of plants were recorded at interval.

## 3. RESULTS AND DISCUSSION

### 3.1. Collection and characterisation of tannery effluent

The tannery effluent was collected from local industries around Vaniyambadi District, Tamil Nadu and analyzed for Physicochemical and biological characteristics (Table No 2; Figure 1) (Report attached at the end of results).

**Table No 1:** Statistical data on Effect of waste water on plant growth.

| Treatment | Germination % | Shoot length cm | Root length cm | Seedling length cm | Chlorophyll mg/g |
| --- | --- | --- | --- | --- | --- |
| Filtered effluent | 70.61 ± 2.01 | 52.78 ± 0.98 | 11.42± 0.43 | 64.50± 1.77 | 2.58± 0.08 |
| Waste after nanosynthesis | 91.52± 2.83 | 55 ± 1.47 | 13 ± 0.19 | 69.2± 1.97 | 3.6± 0.13 |
| Bioremediated effluent | 74.77± 1.62 | 48.9 ± 1.05 | 12.52± 0.34 | 59.6± 2.4 | 2.34± 0.07 |
| Untreated tannery effluent | 45.79 ± 1.21 | 21.47± 0.68 | 6.25± 0.14 | 26.25± 1.25 | 2.25± 0.01 |
| Tap water | 89.3 ± 2.28 | 38.3± 1.06 | 12 ± 0.12 | 49.97± 1.28 | 2.4± 0.04 |

**Table No. 2.**
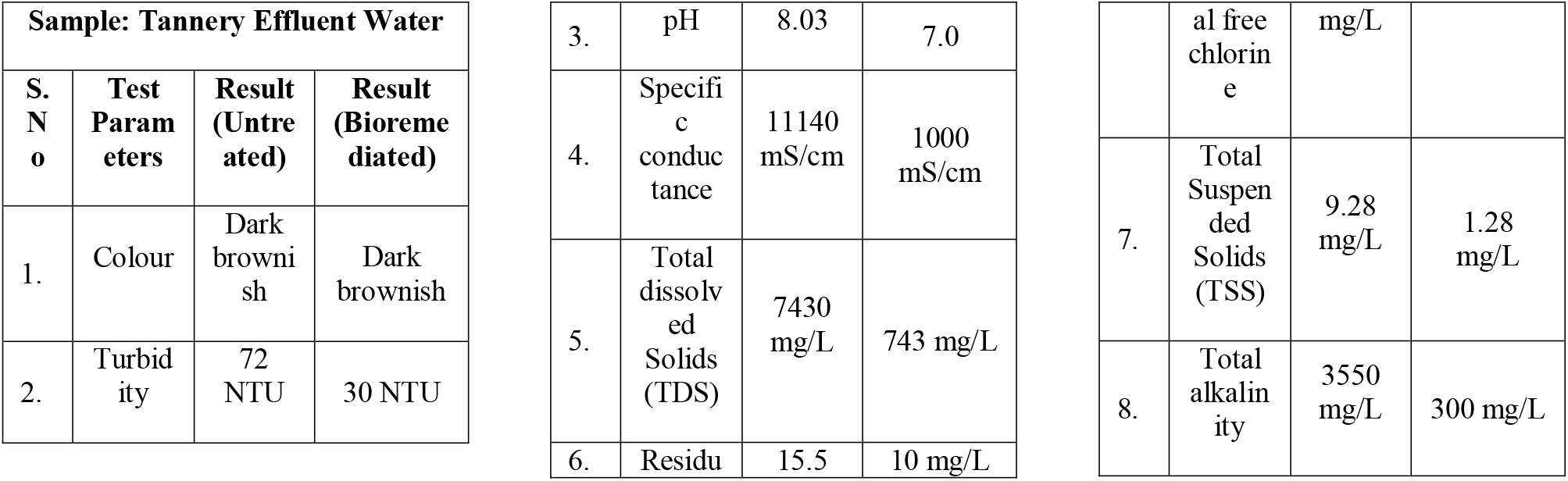
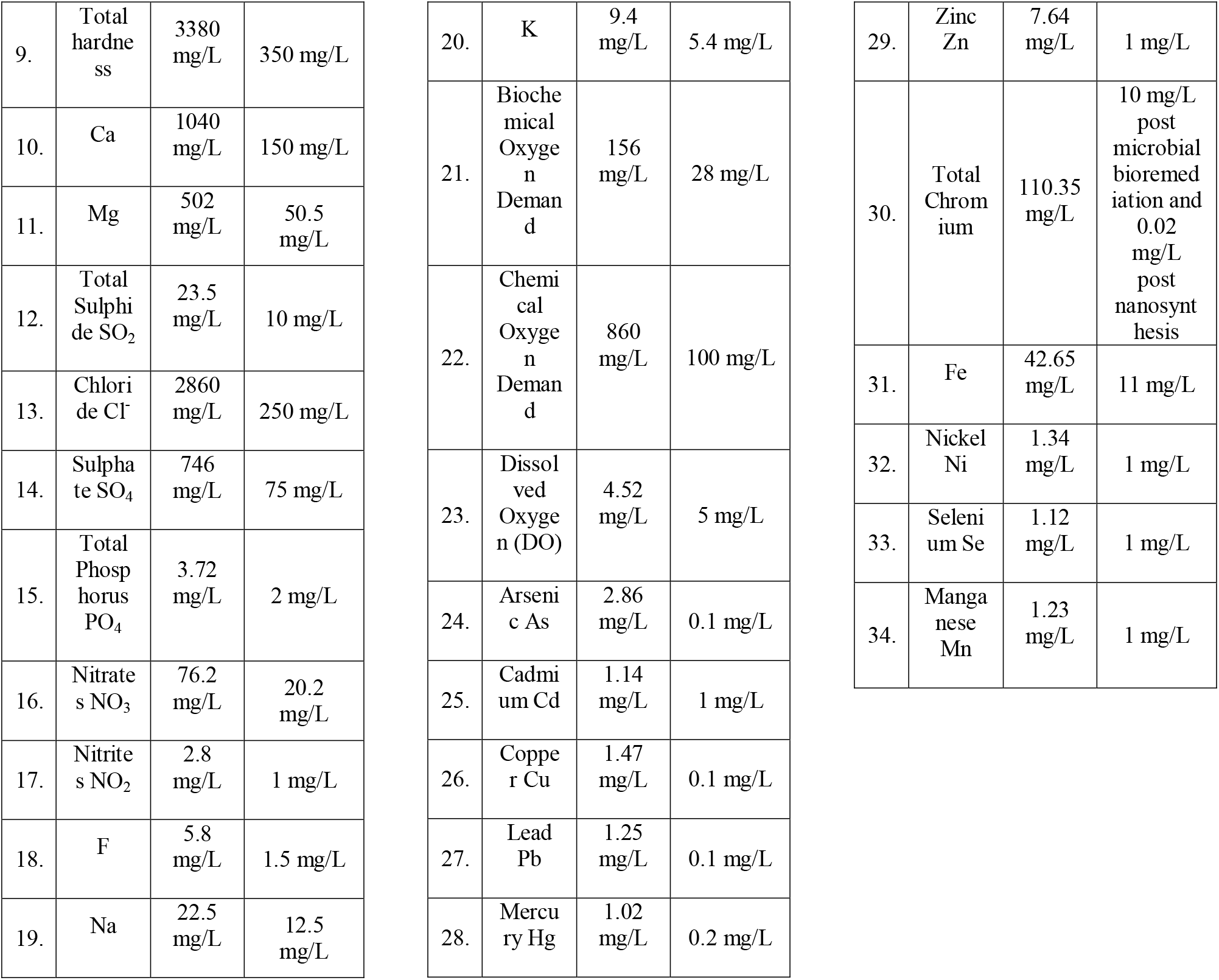
Analysis of Physical and Chemical parameters of untreated and treated tannery effluent.

**Figure No 1.**
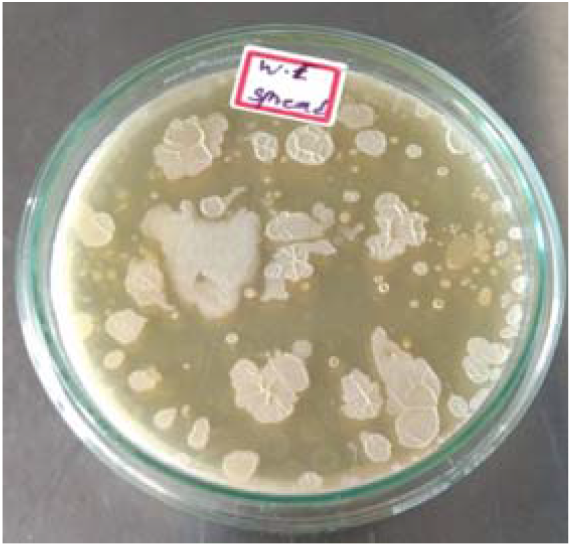
Spread plate of tannery effluent to study the microbial count.

### 3.2. Isolation of Cr (VI) reducing bacteria and tolerance study

Among 50 isolates, only three namely BT1, BT2, M11 showed tolerance to Chromium at a concentration of 100µg/ml. The tolerant isolate identified as *Bacillus sp* was sub-cultured and maintained for further studies. Khookhae et al (2022) have investigated indigenous bacteria isolated from tannery effluents and tested for tolerance to Chromium (VI). Similarly, a previous study by Kabir et al., (2018) have also discussed on isolation of bacteria capable of chromium (VI) reduction from tannery effluent and were characterized to understand the taxonomy of the chromium reducing bacteria. Sanjay et al., (2020) have focused on the isolation of chromium reducing bacteria from tannery industry effluents. They have reported two significant isolates of *Klebsiella pneumoniae* and *Mangrovibacter yixingensis*.

### 3.3. Bioremediation of Chromium and estimation using spectroscopic method

There observed visual colour changes after 48 hour incubation with potent isolate (Figure No 2). The chromium VI was reduced to Chromium III upon bioremediation using potent isolate *Bacillus subtilis* (Table No 3). The concentration of Cr VI was estimated using 1,5 diphenylcarbazide or DPC method. The total chromium was analyzed from Omegaa laboratories, Namakkal as 110.5 mg/L. After nanosynthesis, the Cr (VI) was measured to be 0.02-0.03 mg/L. There has been approx. 99% of Chromium conversion to nanoparticles and this was supported by other characterization studies. The treatment of effluent using microbes reduced the total Chromium to 10mg/L from 110.35 mg/L which is around 90% removal. Further reduced to 0.02 mg/L total Chromium as estimated using DPC method after nanoparticle synthesis and solid separation (Table no 3). Chromium loads were significantly reduced by the treatment train Total Cr was reduced significantly as 110.35 to 10.00 mg/L in bioremediation (90.94%) and 0.01-0.05 mg/L in nanoparticle synthesis (99.95-99.99%) of total Cr. Against regulatory standards, the bioremediated effluent was not within the regulatory limit of CPCB tannery-effluent discharge of total chromium (2 mg/L), but the nanosynthesized effluent was in full compliance. On comparison with potable-water standards, the nanosynthesized effluent at 0.01 mg/l and 0.05 mg/l are below the WHO/BIS total chromium concentration (0.05 mg/L) and meet the recommendation value, respectively. These findings validate the primary findings that: where water quality improvements are a priority, bioremediation alone can reduce chromium sufficiently, but nanosynthesis will be necessary to achieve discharge compliance and drinking-water guideline levels. There are reports on the bioremediation of chromium and arsenic by microalgae in a recent study by Ganguly et al., (2024). Similarly, Seragadam et al., (2021) have mentioned the reduction of hexavalent chromium using *Bacillus* sp. Similarly, Pattnaik et al., (2022) have reported microbial bioremediation of hexavalent chromium by several species of *Bacillus, Staphylococcus, Micrococcus, Acinetobacter, Cellulomonas, Escherichia, Serratia, Pseudomonas, Klebsiella, Achromobacter and Ochrobactrum*.

**Figure No 2:**
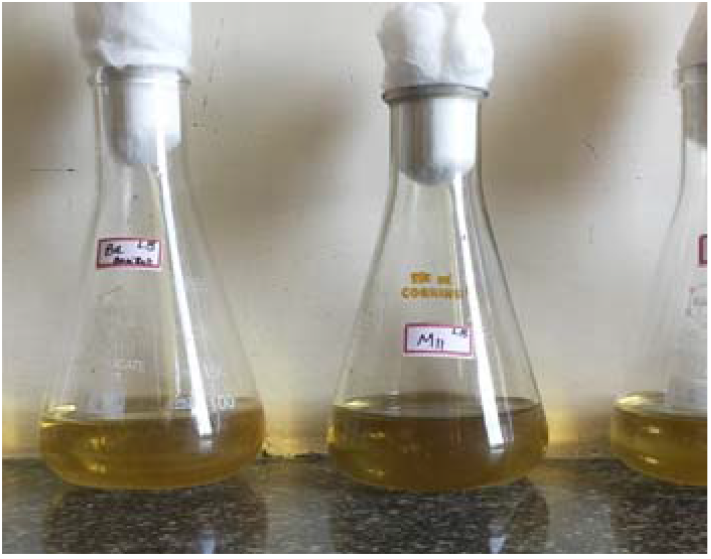
Potent isolate showing Chromium tolerance BT 1 the most tolerant.

**Table No. 3.** Rate of Chromium removal(mg/L) post treatments.

**Table No. 2** Analysis of Physical and Chemical parameters of untreated and treated tannery effluent **Table No. 3** Rate of Chromium removal(mg/L) post treatments
| Stage | Total Cr (mg·L<sup>-1</sup>) | % removal compared to initial concentration |
| --- | --- | --- |
| Effluent (initial total Cr) | **110** | 0.00% |
| After bioremediation (microbial) | **10** | **90.91%** |
| After NP synthesis (final using *Tridax*) | **0.02** | **99.98%** |
| % Recovery (dissolved + solids) | **100.00%** | — |

### 3.4. Molecular characterization of hexavalent chromium reducing bacteria

Among the isolated bacteria, two strains namely BT1/M11 and BT2 were found to have significant ability for Chromium (VI) bioremediation (Figure No 1 and 2). However, BT1 had stable bioremediation capacity and was used further for the bioremediation. The strain BT1 was identified as *Bacillus subtilis* and strain BT2 was identified as *Enterobacter* sp. The 16s rRNA sequencing was done for BT1 and the accession number was obtained from NCBI GenBank-PQ764883. There are reports of several species of microorganisms that help in bioremediation of hexavalent chromium. Ghosh et al., (2021) have highlighted in their review on different microbes helping in the reduction of hexavalent chromium like algae, fungi, bacteria and plant species.

### 3.5. Production and Characterisation of Chromium oxide nanoparticles using *Tridax procumbens* (Green synthesis)

The nanoparticles produced had a greenish tint and the filtrates were dried and send for FSEM analysis, Zeta potential analysis and XRD analysis. Green nanosynthesis is widely researched topic for its efficiency and eco friendliness (Figure no 3). There are recent reports of green nanosynthesis of manganese di oxide using *Tridax procumbens* for dye degradation and antimicrobial activity (Veena et al., 2024). Ghotekar et al., (2021) have reported the production of chromium nanoparticles using *Tridax procumbens* leaves and reported several biomedical applications on it. Yasmeen et al., (2023) have reported the use of several analyses for studying the characteristics of nanoparticles synthesize from Cassia fistula by green nanosynthesis method. The analysis included SEM, TEM, AFM, FTIR, Uv-Vis spectroscopy, DSC, Raman spectroscopy along with analysis of their electrochemical properties.

**Figure No 3:**
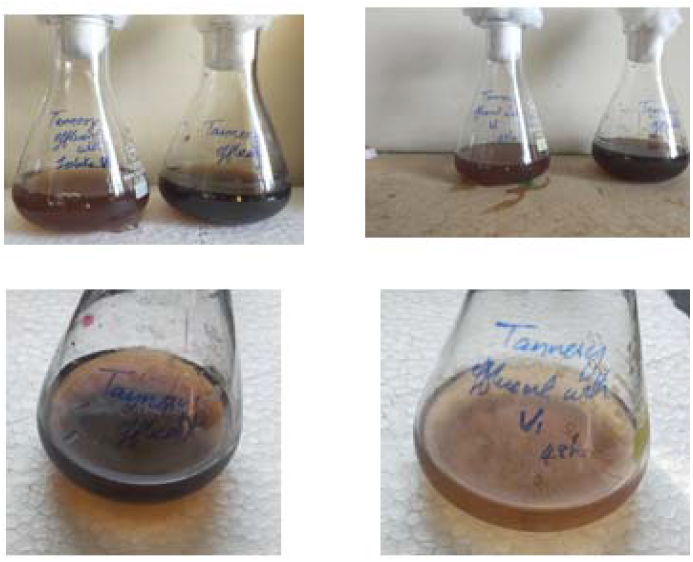
Bioremediation of Chromium (VI) using M11/BT1. Best result after 48 hour.

#### 3.5.1. XRD analysis

The X ray diffraction pattern of the nanoparticle shows sharp pure peaks appearing at different values of 2θ at 24°, 28°, 32°, 33°, 36°, 41°, 50°, 54°, 63°, 65°, 72° and 76° showing crystallinity of the nanoparticle. The mean crystalline size is 12.33nm (Figure no 4,5).

**Figure No 4:**
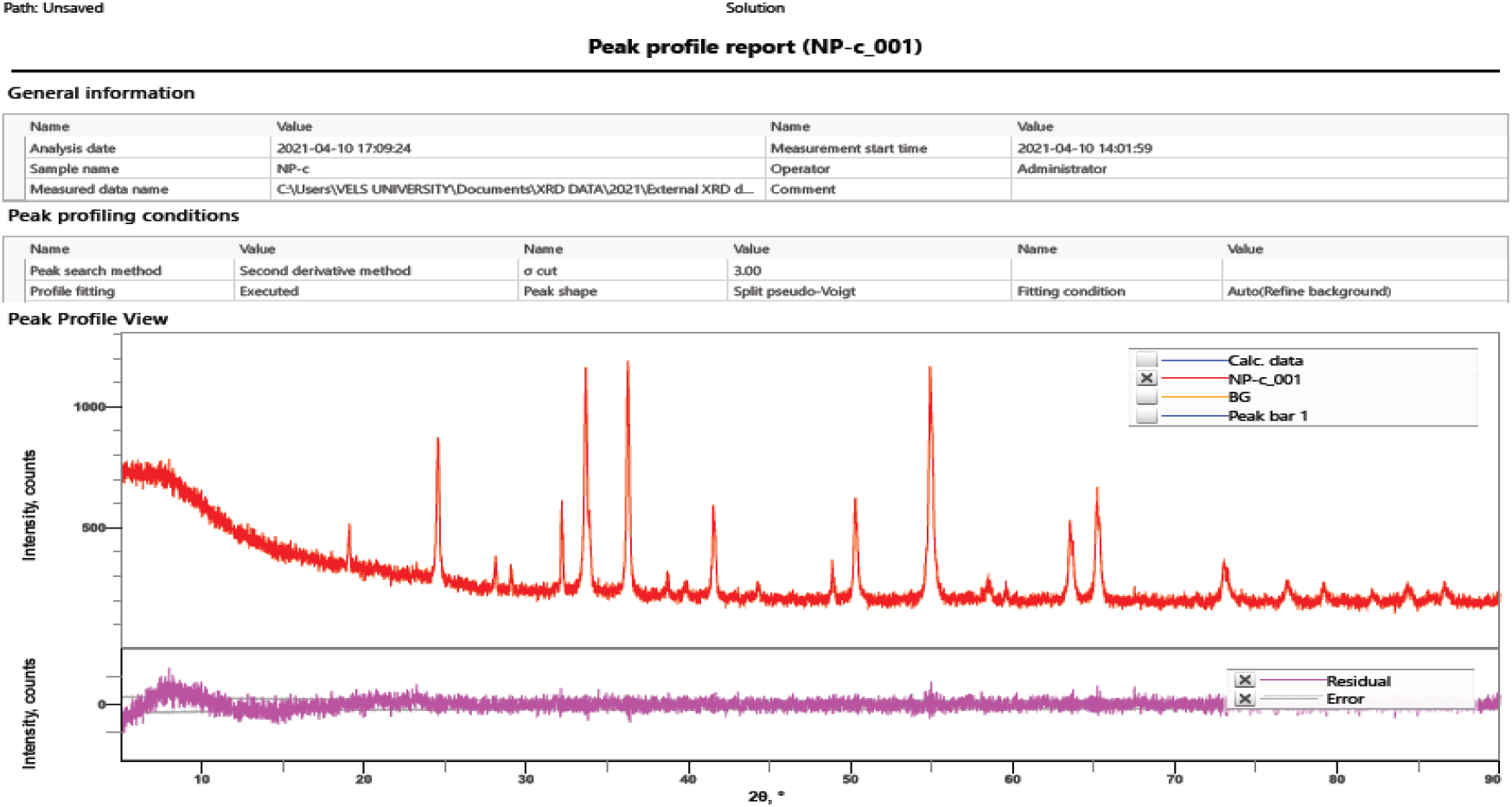
XRD analysis

**Figure No 5:**
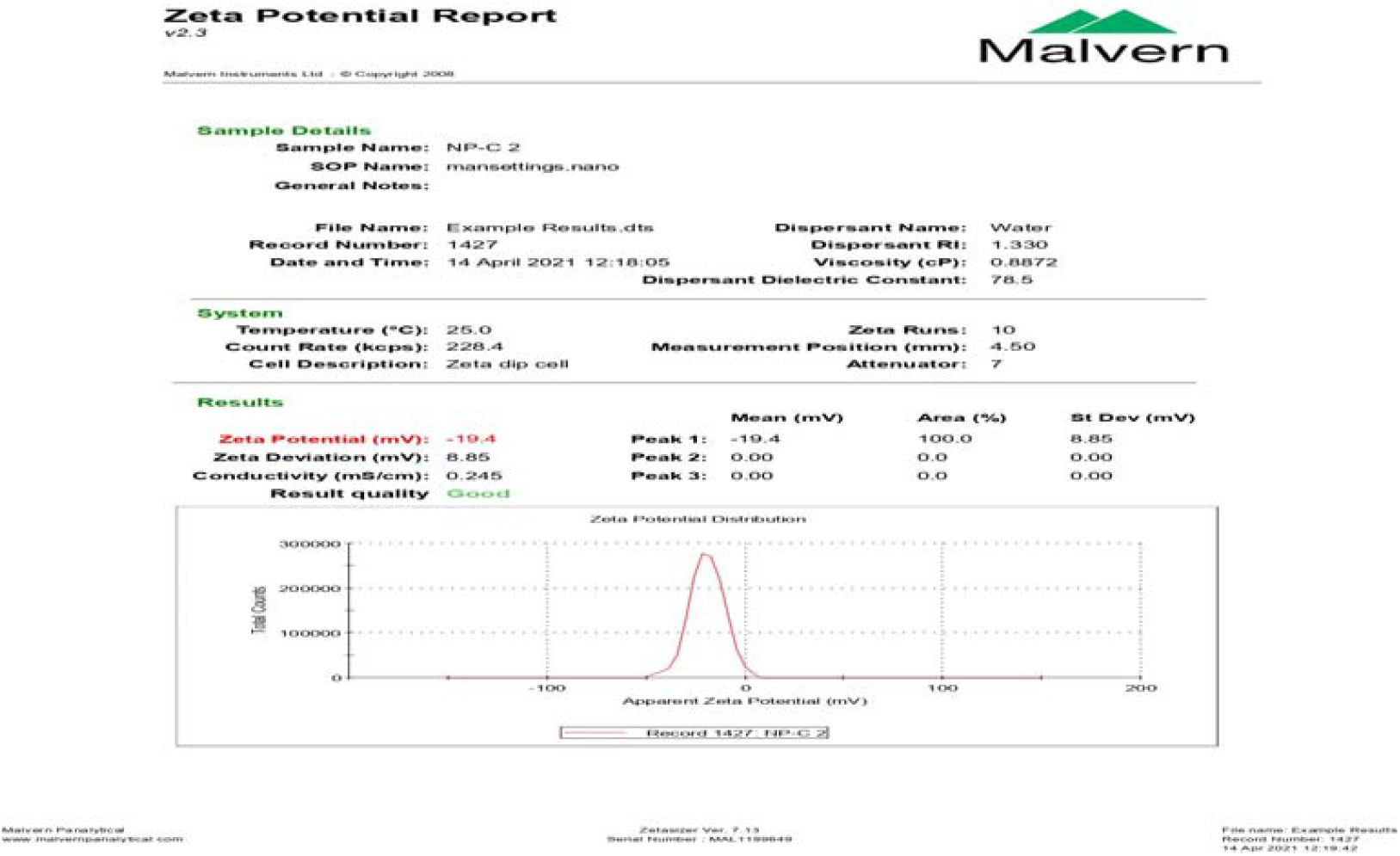
Zeta potential Report

**Figure No 6:**
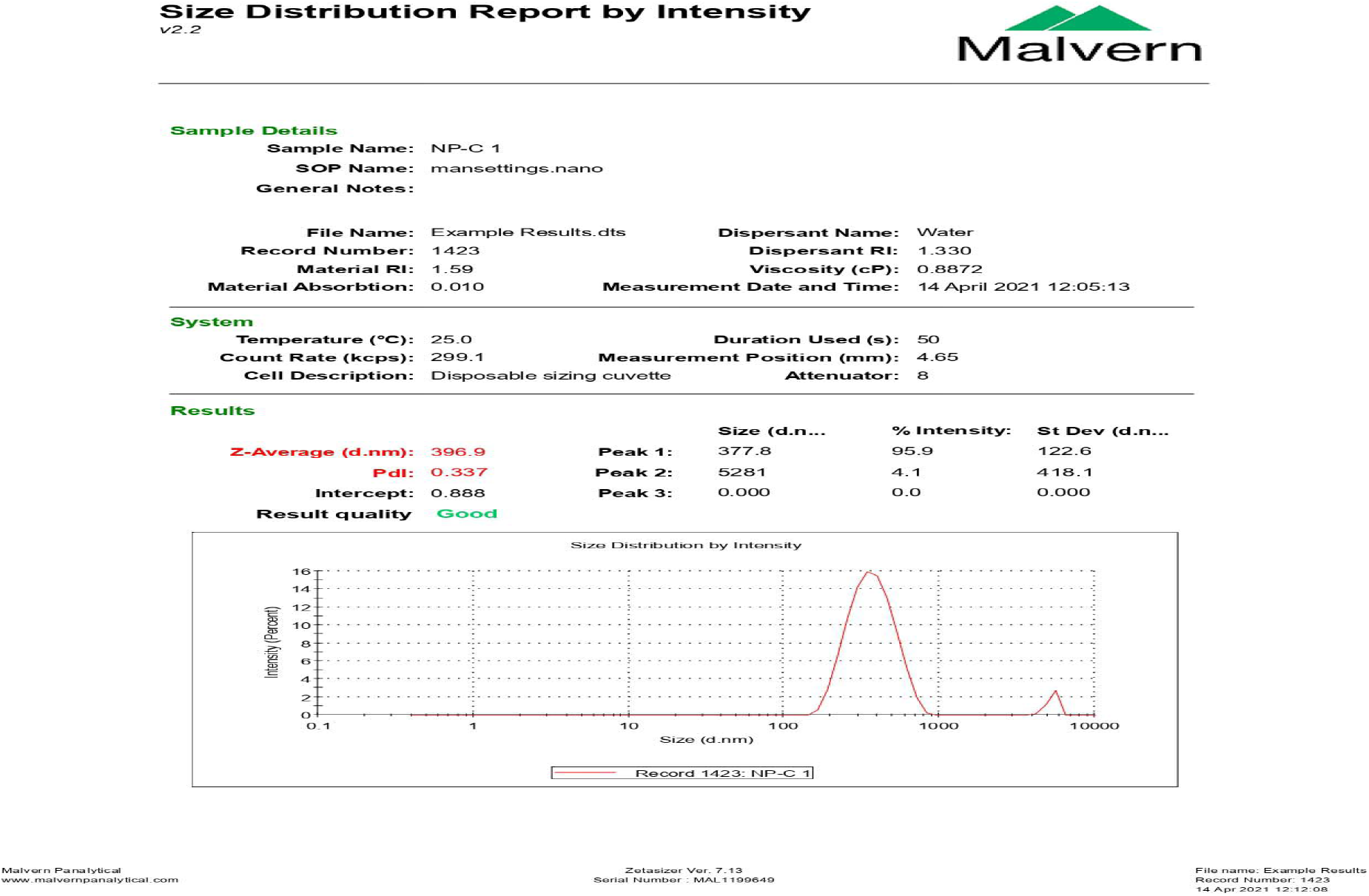
Size distribution –Zeta potential report

There are previous reports by Hussain et al (2020) and Rheima et al (2020) which focussed on the XRD an analysis of Chromium oxide nanoparticle. Marjani et al., 2021 have analyzed chromium nanoparticle and reported the crystalline size to be 12.65 nm. Ghotekar et al., (2021) have also reported on crystallographic characterization using XRD and have also discussed the shape and size of nanoparticles obtained from microbial synthesis like hexagonal shaped 36 nm nanoparticle from *Aspergillus niger* biosynthesis and spherical 50 nm shaped NPs from *Bacillus subtilis*.

#### 3.5.2. FSEM analysis

The surface morphology of synthesized Chromium oxide was studied using SEM analysis. The synthesized nanoparticles were observed as fine particles of greenish colour (Figure no 7,8). Cr2O3 NPs are having particle size in the range of 20 nm – 70 nm. The size distribution histograms for the respective nanoparticle provided their respective sizes as 32.9 ± 13.3 nm, 32.2 ± 9.3 nm, 31.4 ± 9.5 nm, 28.8 ± 8.9 nm, respectively. These findings were in accordance with earlier reports where LS has been widely used to characterize nanoparticle dispersion and aggregation behavior (Ghotekar et al., 2021)

**Figure No 7:**
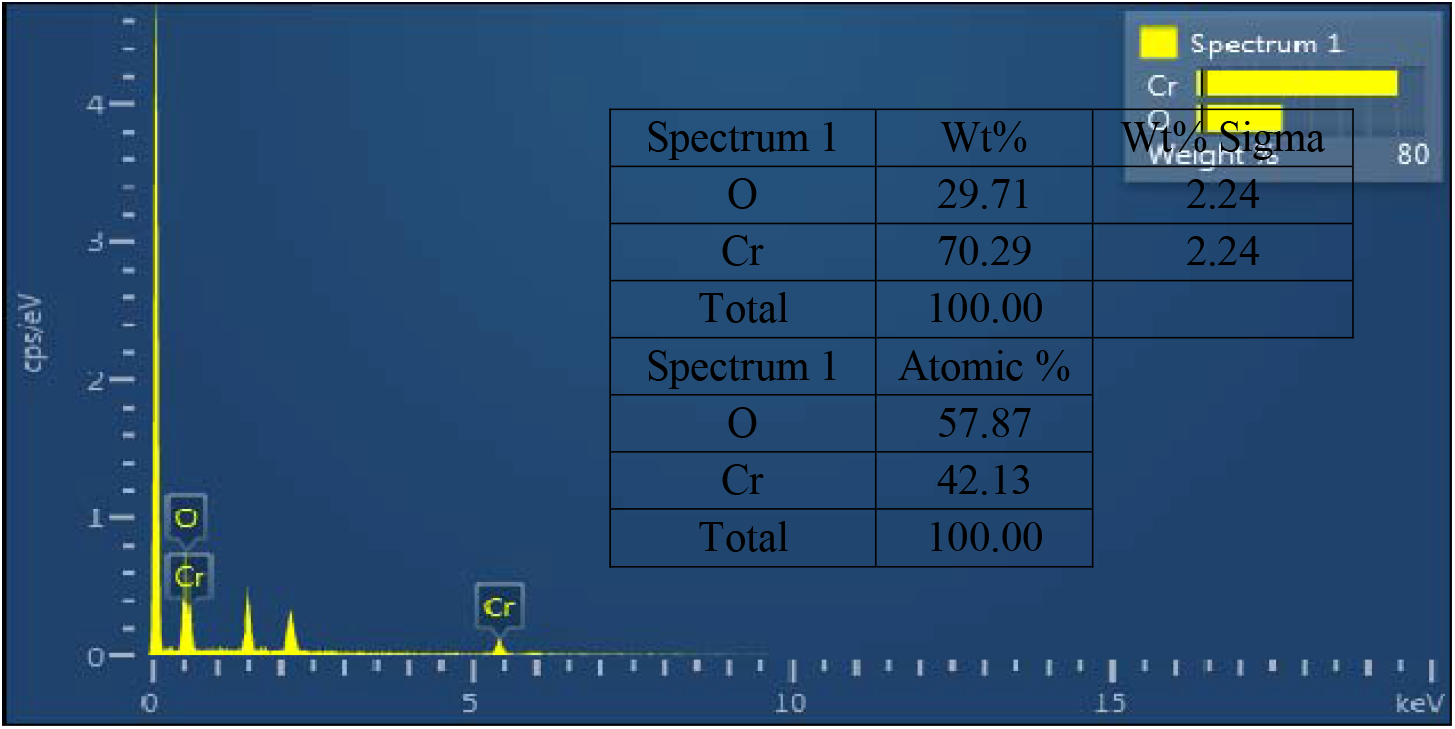
EDAX report

**Figure No 8:**
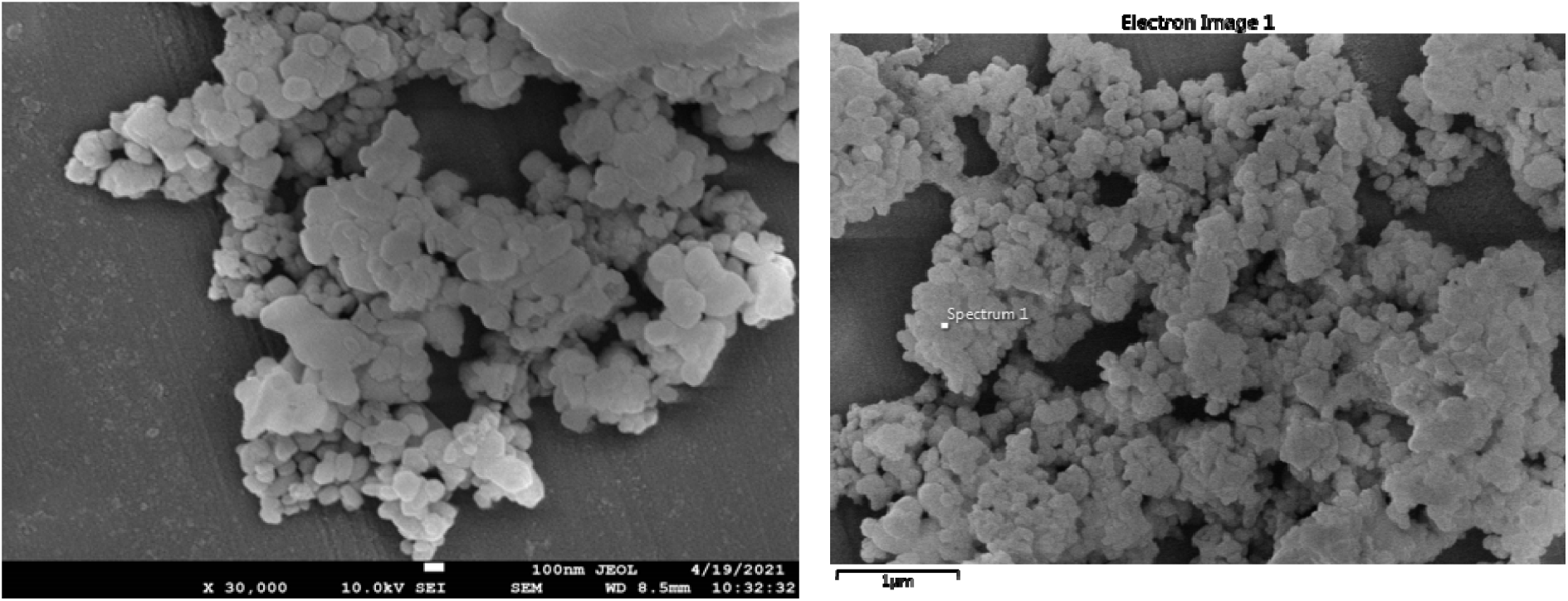
FSEM report (x 3000)

#### 3.5.3. Zeta potential analysis/ Particle size

Zeta potential and particle size was observed for the synthesized chromium nanoparticles. The charge of the nanoparticle was −19.4 mV (Figure no 6). The size distribuition Z average was about 396.9 nm. Ghotekar et al., (2021) have also reported on DLS analysis of nanoparticles to study the size distribution and agglomeration of nanoparticles.

### 3.6. Seed Germination

Green gram plant was grown with bio-remediated tannery effluent and observed for growth. The experiment was performed both in pot and field. Total of four set of experiment was performed namely growth of green gram grown using filtered effluent, using waste effluent after nanosynthesis, using bioremediated effluent and untreated tannery effluent (Figure no 9). It was observed that growth was at its best when waste effluent after nanosynthesis was used for cultivation in both pot and field experiment. There are several reports and research on the effect of seed germination using several industrial effluents. It was observed that the highest germination percentage was consistently observed in seeds treated with waste effluent after nanosynthesis of chromium oxide (Table No 1). This suggested that nanosynthesis not only reduced the toxicity of the effluent but has shown the nutrient profile that may have promoted better seedling establishment. One way ANOVA analysis was conducted to understand the treatment effects and null hypothesis. However, a bioremediated effluent with zero chromium and its application for seed germination may be innovative and have not yet been reported widely. Mythili et al., (2011) have reported the germination of black gram and sunflower using bioremediated tannery effluent. Similarly, Hailu et al., (2019) have also studied on effect of tannery effluent on seed germination of maize varieties.

**Figure No 9:**
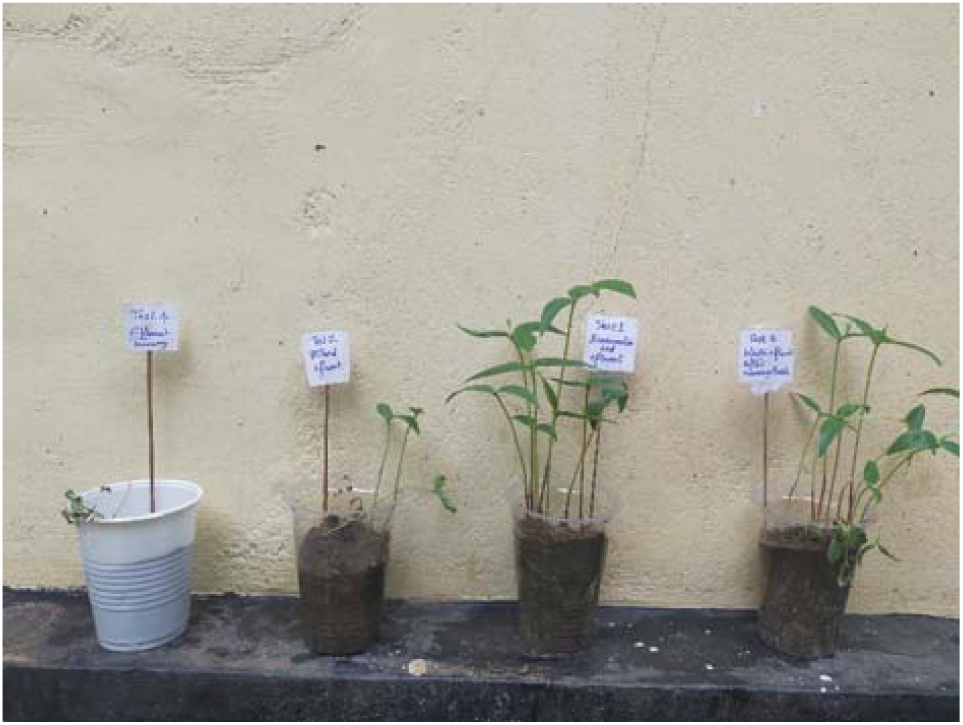
Seed germination experiment-pot (21 days) Test 1 - Bioremediated effluent; Test 2 - Filtered effluent; Test 3 - Waste effluent after nanosynthesis; Test 4 - Tannery effluent

**Figure No 10:**
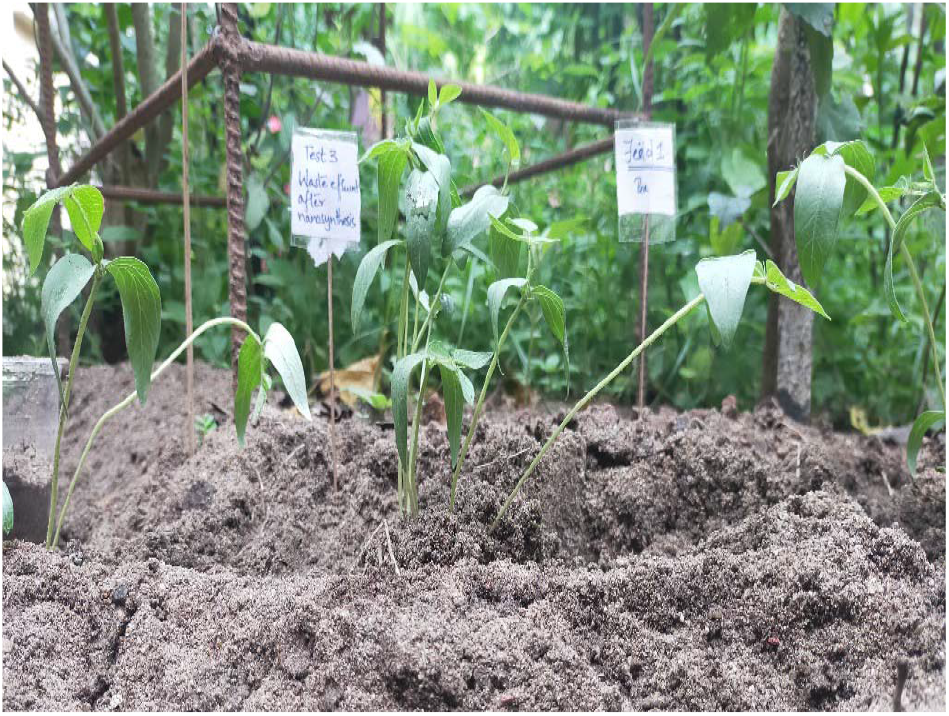
Field Data of Seed germination experiment using Waste effluent after nanosynthesis (21 days)

### 3.7. Statistical Analysis

A one-factor analysis of variance has shown that there is a significant difference between the categorical variable Treatment and the variable. The following was observed-Value F = 186.85 for germination %, F = 487.3 for shoot length, F = 275.33 for root length, F = 229.99 for seedling length and F = 129 for chlorophyll content with p = <.001 for all variables. The data exhibited strong treatment effects. Thus, with the available data, the null hypothesis is rejected. The rejection of the null hypothesis confirms that the variation observed was not due to chance, but rather to the nature of the effluent applied.

## 4. CONCLUSION

Vellore district of Tamil Nadu houses several leather industries, both large and small scale industries which release tonnes of waste into the environment causing severe menace to agriculture and groundwater systems. Most of the unit lac treatment plants due to high cost for plant establishment. The current projects aim to bio-remediate the fatal heavy metals in the tannery effluent using natural microbes and use the treated effluent for agriculture thus recycling the effluent cost effectively. The project due to its cost effectiveness can be implemented by the small-scale industries at the same time promoted to agriculture purposes.

## Acknowledgements

The author would like to acknowledge Sacred Heart College (Autonomous), Tirupathur, Tamil Nadu, India for providing research grant-Don Bosco Research Grant as financial aid. I would like to acknowledge Lovely Professional University, Punjab, India and IISc, Bangalore for their instrumentation facilities.

## Author contribution

Dr Neethu Asokan-concept and idea, experimental work and research, data analysis and compilation, writing of paper, editing and revision.

