## Supplementary File for "SUSTAINABLE MICROBIAL BIOTRANSFORMATION OF CR(VI) TO CR(III) IN TANNERY EFFLUENT AND ITS VALORIZATION INTO CR(III) NANOPARTICLES VIA *TRIDAX PROCUMBENS*-MEDIATED GREEN SYNTHESIS"

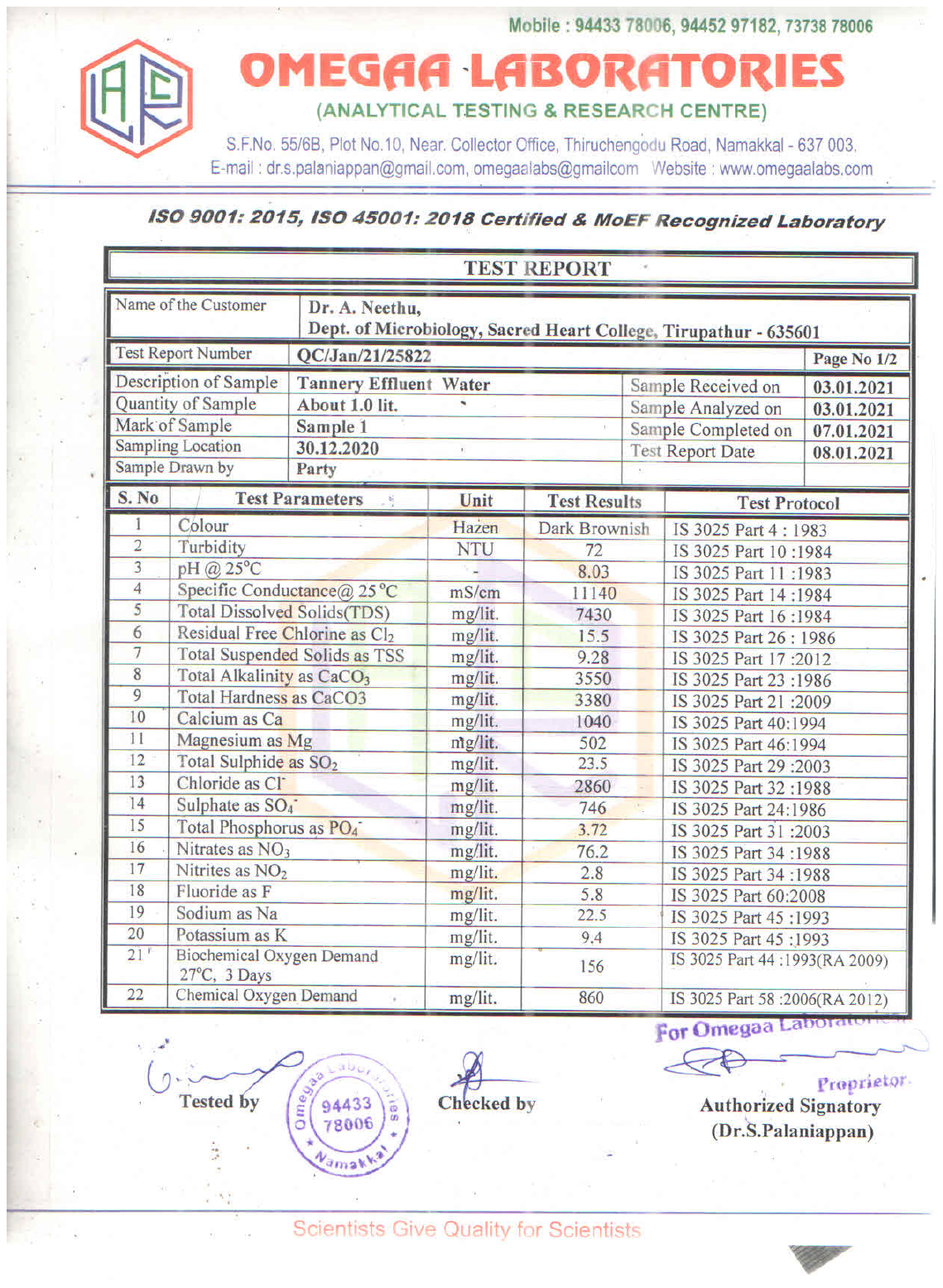

Table No 1 Physical parameter analysis of tannery effluent.

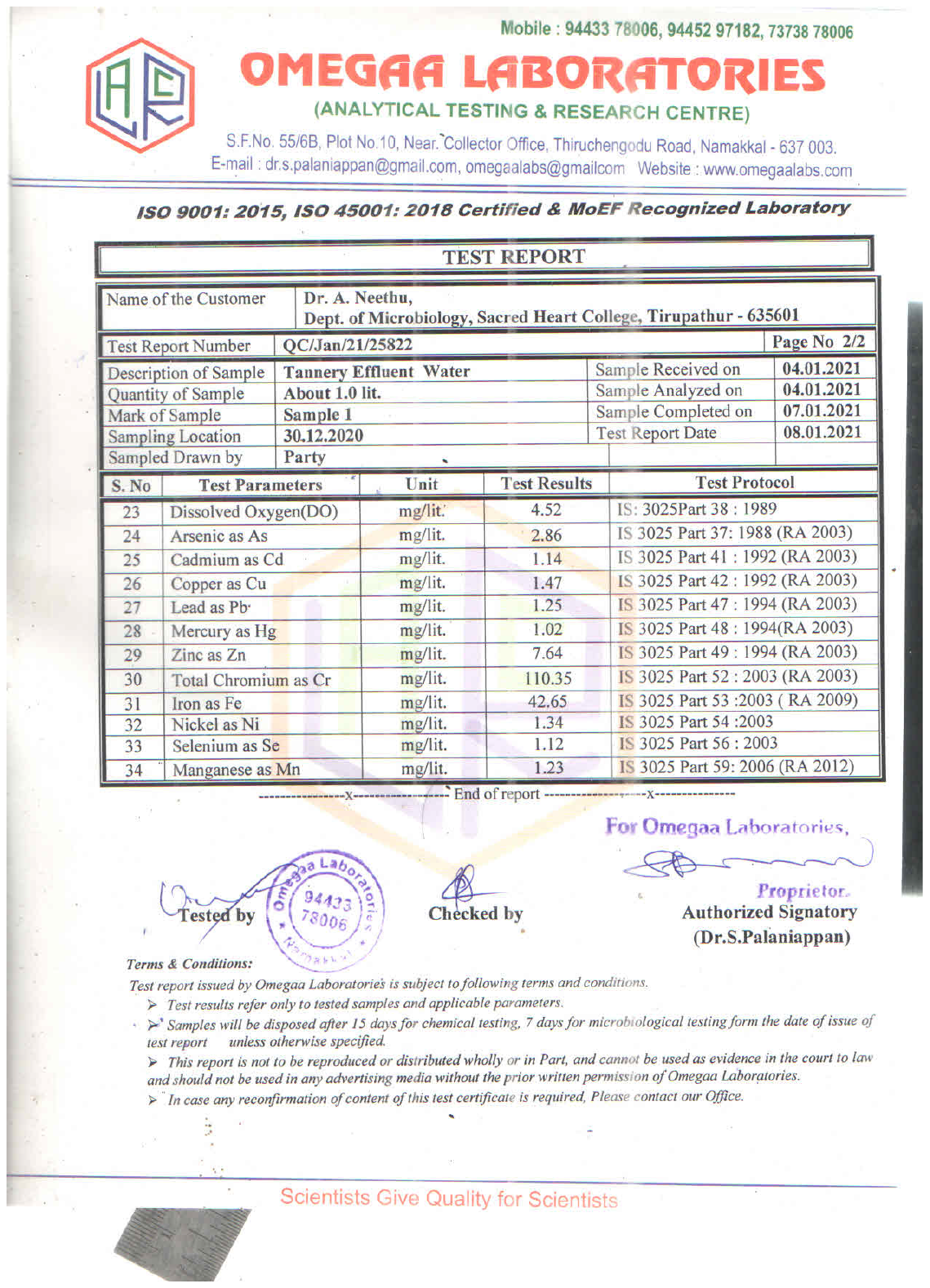

Table No 2 Chemical analysis of tannery effluent.

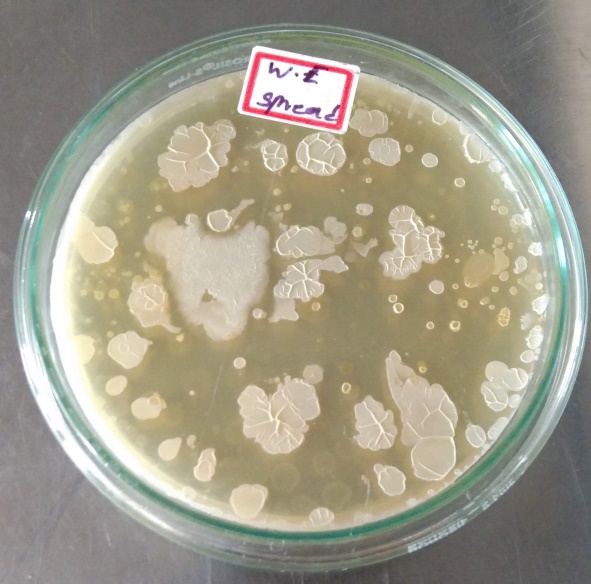

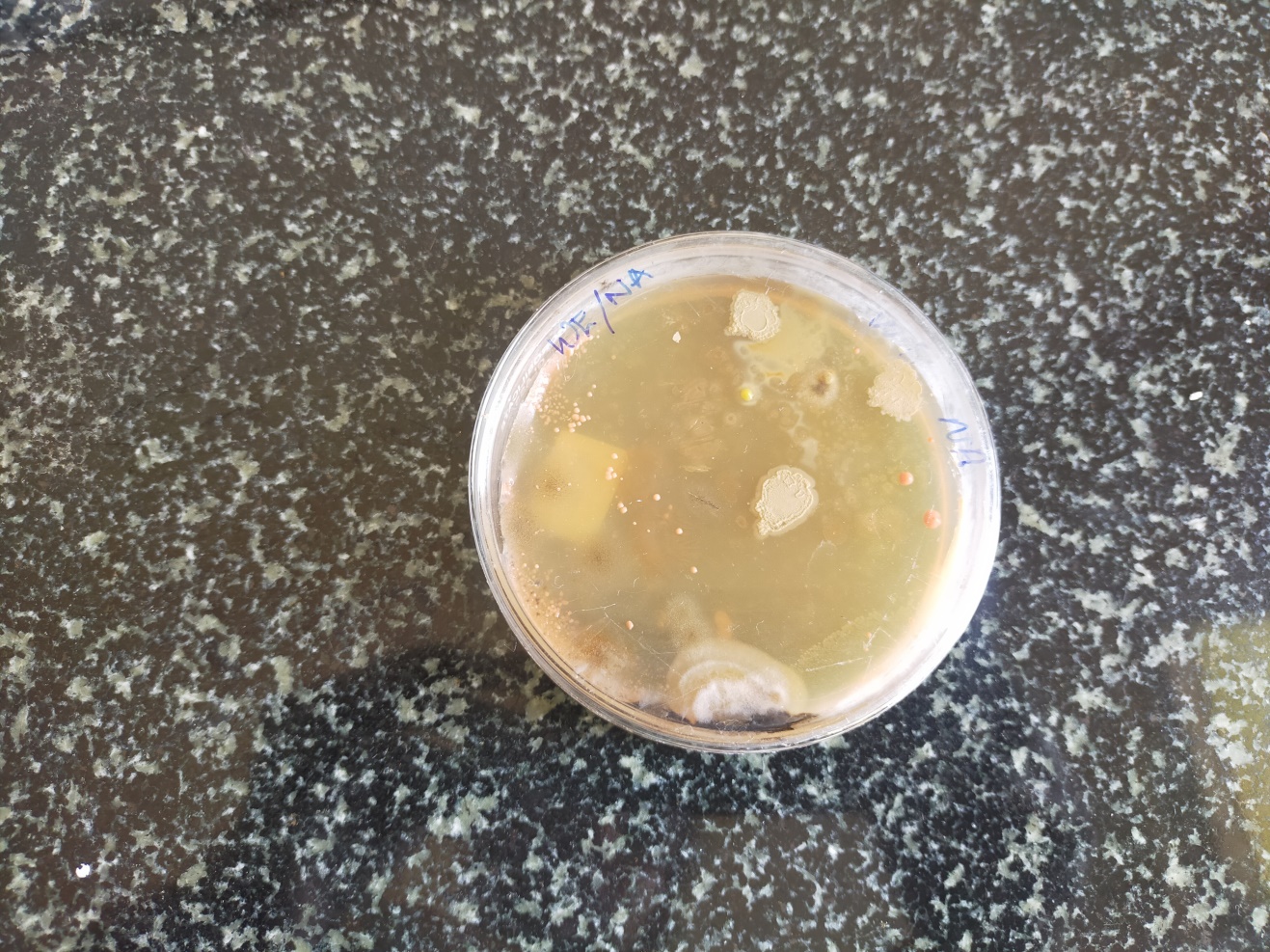

Figure No 1 Spread plate of tannery effluent to study the microbial count.

| **SL No** | **Sample code** | **Colony code** | **Colony morphology** | **Grams Staining** |
| --- | --- | --- | --- | --- |
| 1. | Effluent contaminated soil around Vaniyampadi local leather industry | V1 | Round, Regular, Creamy | G- Bacilli |
| 2. |  | V2 | Irregular , off white |  |
| 3. |  | V3 | Round, Regular, Creamy |  |
| 4. |  | V4 | Irregular , off white |  |
| 5. |  | V5 | Irregular , off white |  |
| 6. |  | V6 | Round, Cream, Raised |  |
| 7. |  | V7 | Irregular, Powdery, white | G+ Bacilli |
| 8. |  | V8 | Round, Regular, Creamy | G- Bacilli |
| 9. |  | V9 | Round, Regular, Creamy |  |
| 10. |  | V10 | Irregular , off white |  |
| 11. |  | V11 | Round, Regular, Creamy |  |
| 12. |  | V12 | Irregular , off white |  |
| 13. |  | V13 | Irregular, Powdery, white | G+ Bacilli |
| 14. |  | V14 | Irregular , off white |  |
| 15. | Effluent collected from local leather industry | M1 | Irregular, Powdery, white | G+ Bacilli |
| 16. |  | M2 | Irregular , off white |  |
| 17. |  | M3 | Round, Regular, Creamy | G- Bacilli |
| 18. |  | M4 | Irregular , off white |  |
| 19. |  | M5 | Round, Cream, Raised |  |
| 20. |  | M6 | Irregular, Powdery, white | G+ Bacilli |
| 21. |  | M7 | Round, Cream, Raised |  |
| 22. |  | M8 | Round, Regular, Creamy |  |
| 23. |  | M9 | Round, Regular, Creamy | G- Bacilli |
| 24. |  | M10 | Round, Regular, Creamy | G- Bacilli |
| 25. |  | M11 | Irregular, Powdery, white | G+ Bacilli |
| 26. |  | M12 | Irregular, Powdery, white | G+ Bacilli |
| 27. |  | M13 | Irregular, Powdery, white | G+ Bacilli |
| 28. |  | M14 | Irregular , off white |  |
| 29. |  | M15 | Irregular , off white |  |
| 30. | Soil collected from around Tannery industry in Vaniyampadi | B1 | Round, Regular, Creamy |  |
| 31. |  | B2 | Irregular , off white |  |
| 32. |  | B3 | Round, Regular, Creamy |  |
| 33. |  | B4 | Irregular , off white |  |
| 34. |  | B5 | Round, Regular, Creamy | G- Bacilli |
| 35. |  | B6 | Round, Regular, Creamy |  |
| 36. |  | B7 | Round, Cream, Raised |  |
| 37. |  | B8 | Round, Regular, Creamy |  |
| 38. |  | B9 | Round, Cream, Raised |  |
| 39. |  | B10 | Round, Regular, Creamy | G- Bacilli |
| 40. |  | B11 | Irregular, Powdery, white | G+ Bacilli |
| 41. |  | B12 | Round, Regular, Creamy |  |
| 42. |  | B13 | Round, Regular, Creamy |  |
| 43. |  | B14 | Round, Cream, Raised |  |
| 44. |  | B15 | Round, Regular, Creamy | G- Bacilli |
| 45. |  | B16 | Irregular, Powdery, white | G+ Bacilli |
| 46. |  | B17 | Round, Cream, Raised |  |
| 47. |  | B18 | Irregular, Powdery, white | G+ Bacilli |
| 48. |  | B19 | Round, Regular, Creamy |  |
| 49. |  | B20 | Irregular, Powdery, white | G+ Bacilli |
| 50. |  | B21 | Round, Cream, Raised |  |

Table No 3: Isolation of microorganism from effluent and Cr contaminated soil.

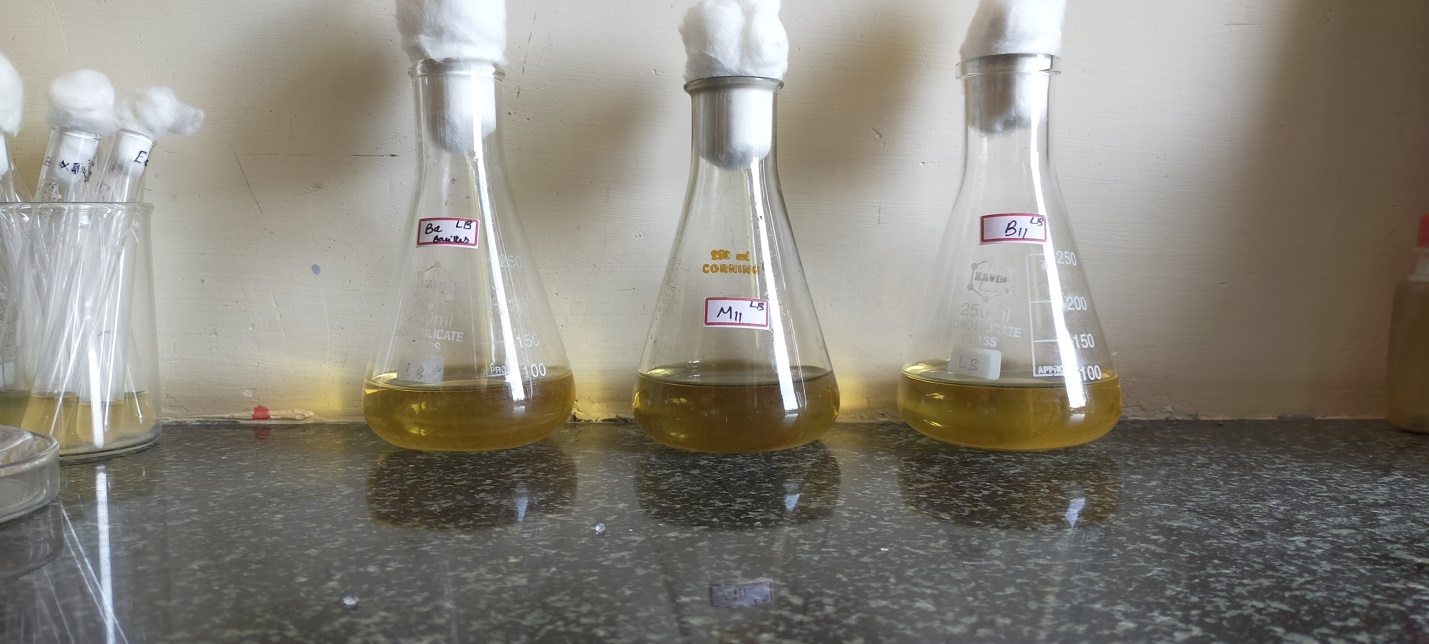

Figure No 2: Potent isolate showing Chromium tolerance M11 the most tolerant.

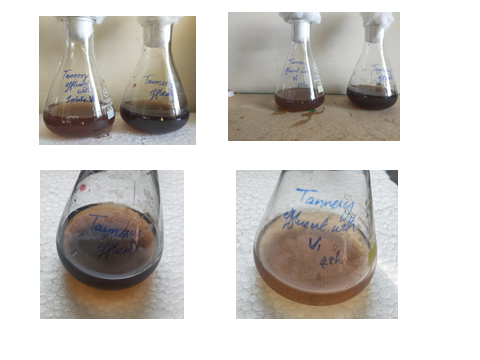

Figure No : Bioremediation of Chromium (VI) using M11. Best result after 48 hour.

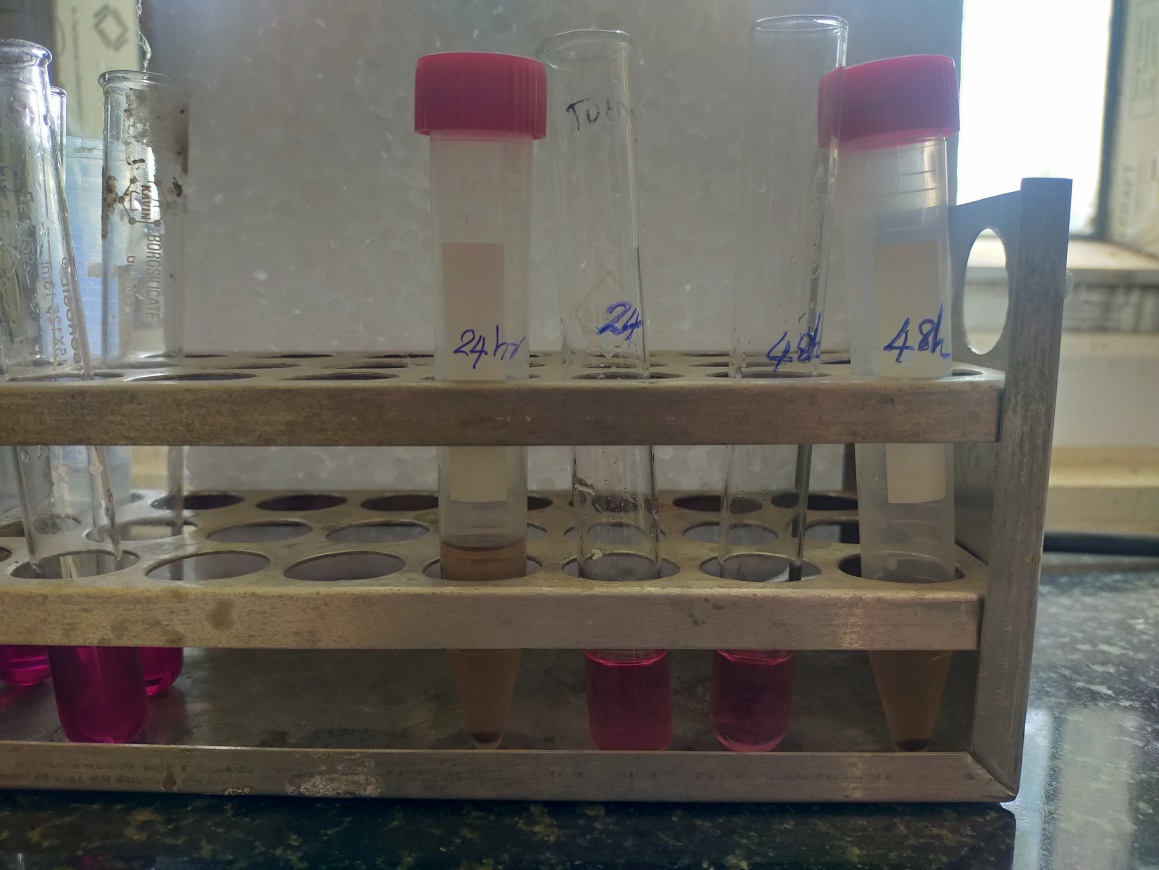

Standard graph - DPC chromium estimation method

| Concentration of Chromium (VI) (mg) | Optical Density at 540 nm |
| --- | --- |
| 0 | 0 |
| 0.2 | 0.12 |
| 0.4 | 0.22 |
| 0.6 | 0.32 |
| 0.8 | 0.43 |
| 1 | 0.54 |

Figure No : DPC Estimation of Chromium after bioremediation with standard graph

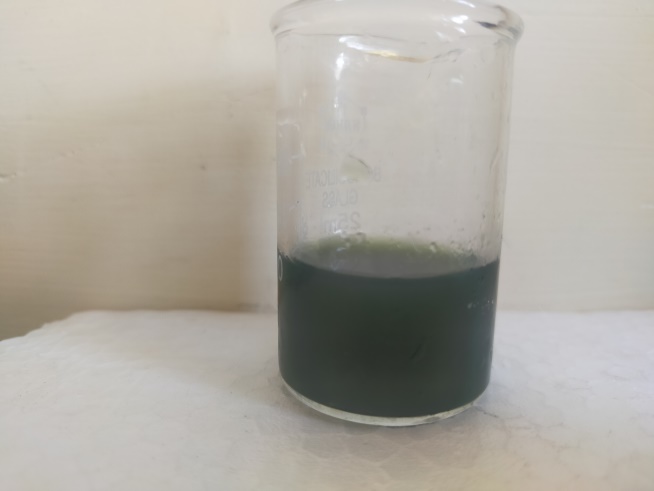

Figure No : Tridax procumbens extract

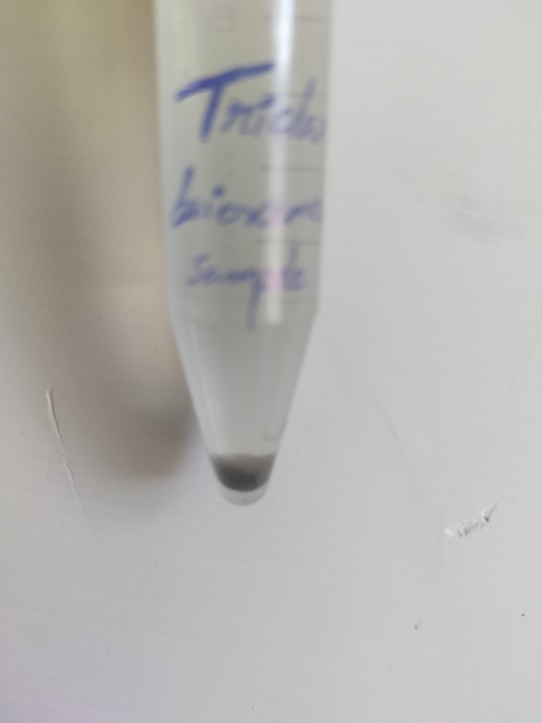

Figure No : Chromium oxide nanoparticle pelletized

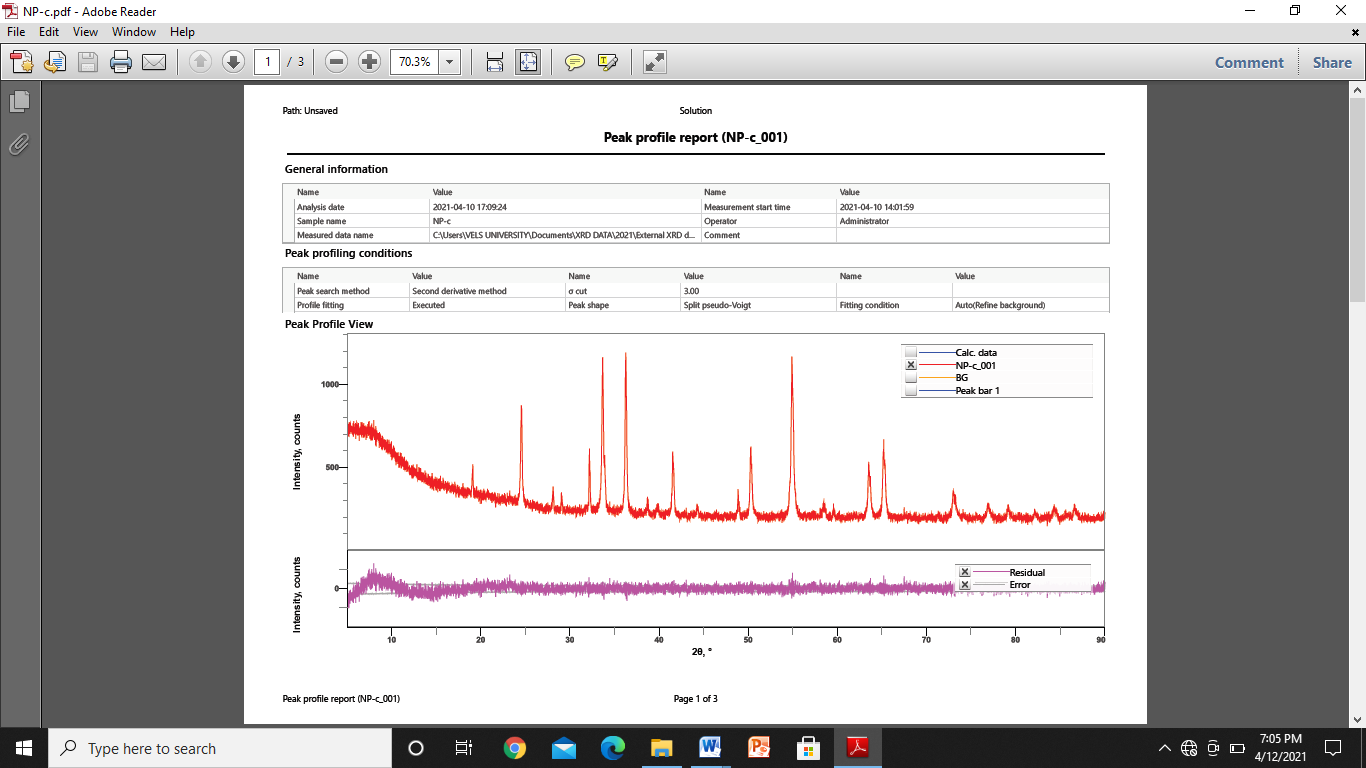

Figure No : XRD analysis

Figure No : Zeta potential Report

Figure No : Size distribution –Zeta potential report

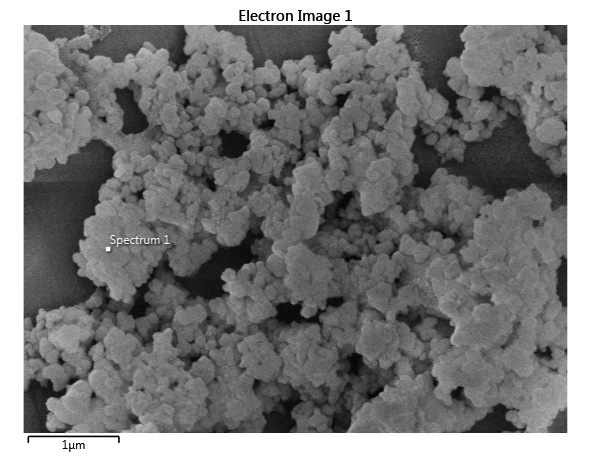

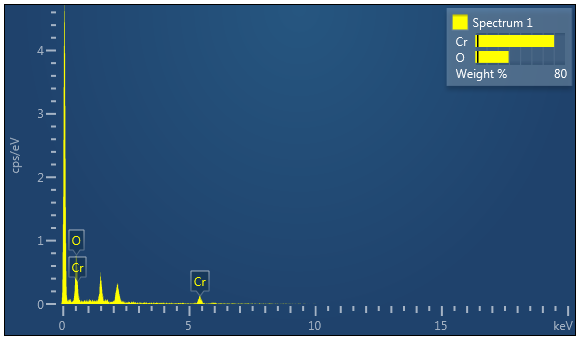

| Spectrum 1 | Wt% | Wt% Sigma |
| --- | --- | --- |
| O | 29.71 | 2.24 |
| Cr | 70.29 | 2.24 |
| Total | 100.00 |  |
| Spectrum 1 | Atomic % | |
| O | 57.87 | |
| Cr | 42.13 | |
| Total | 100.00 | |

Figure No : EDAX report

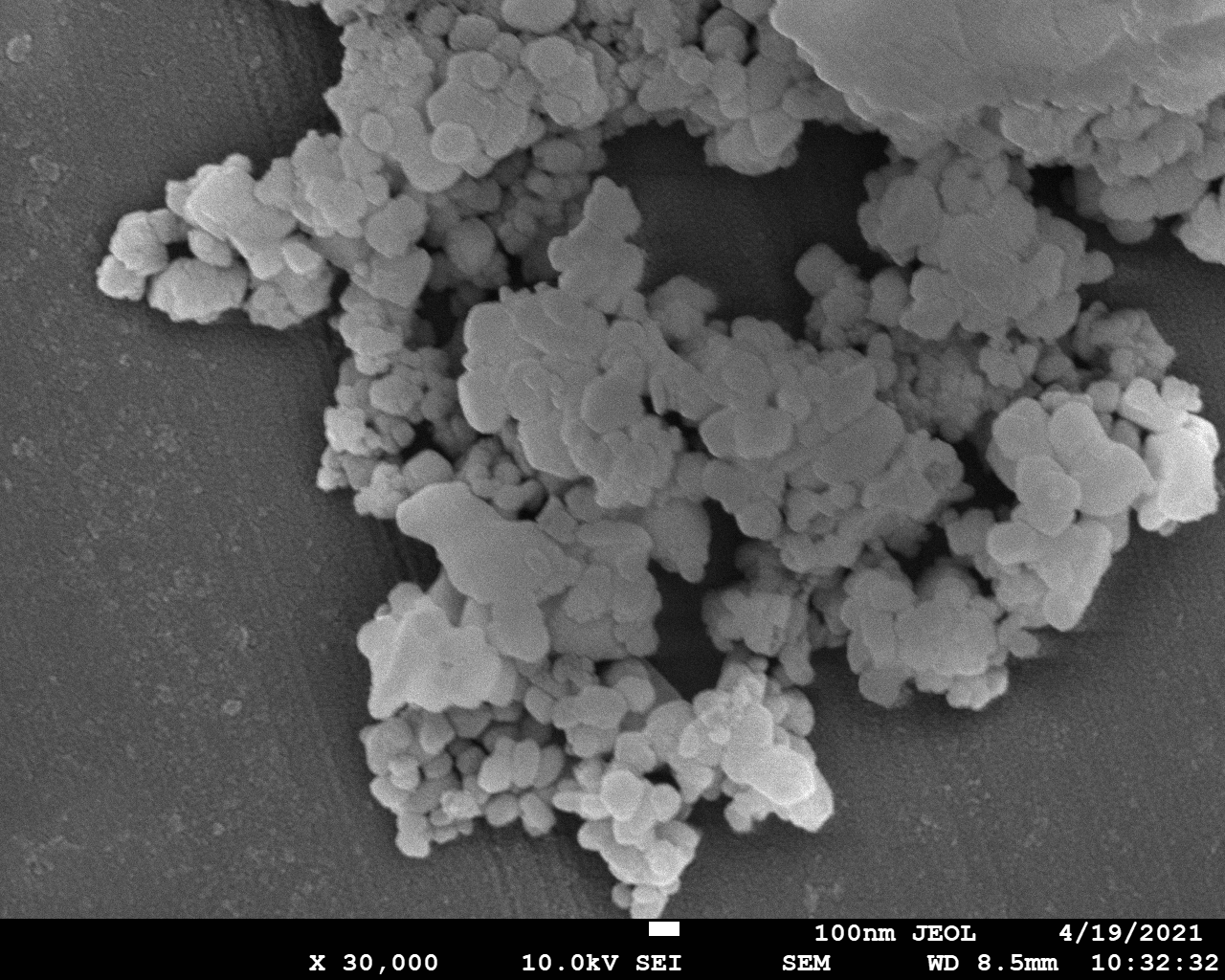

Figure No : FSEM report (x 3000)

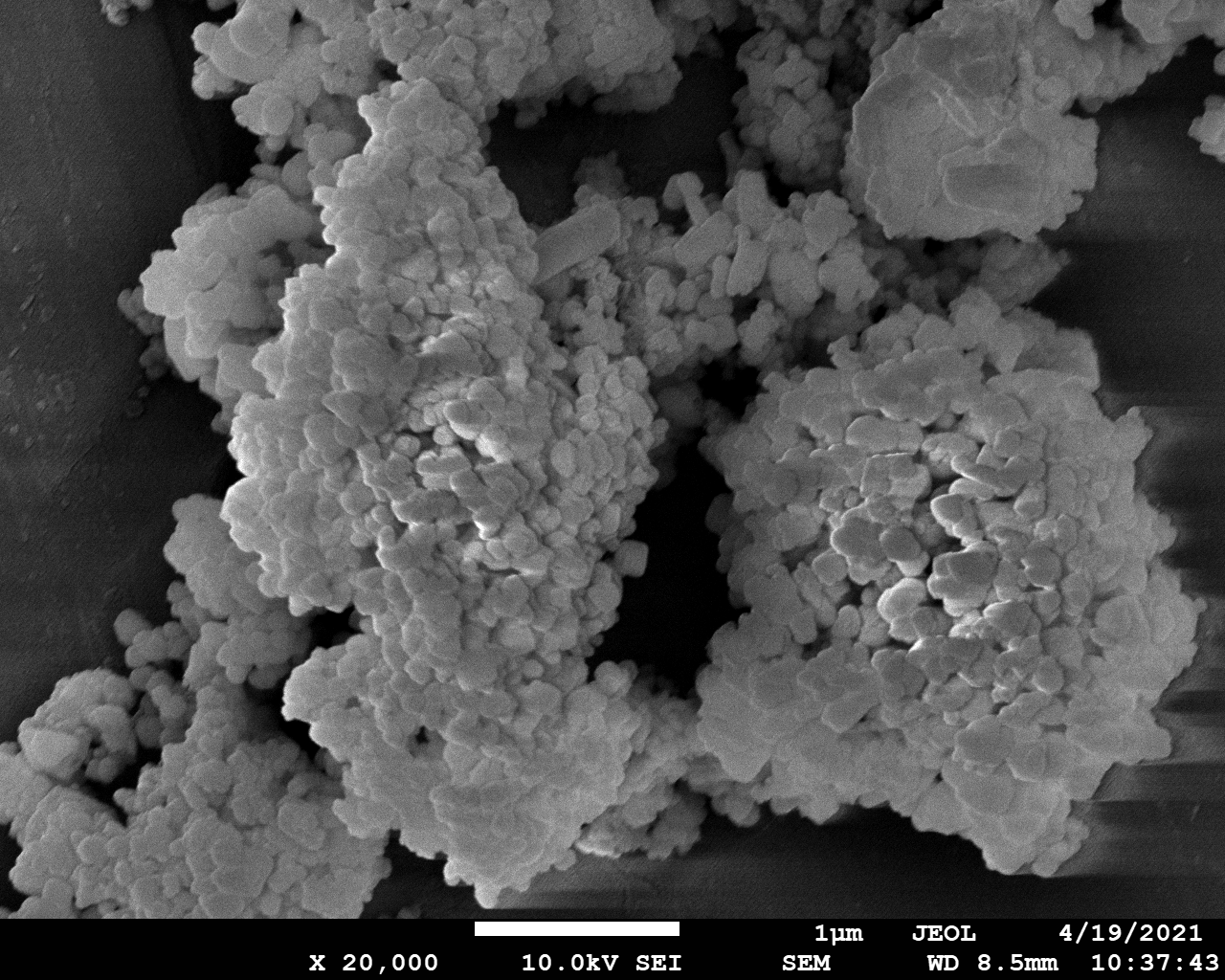

Figure No : FSEM report (x 20000)
